# Taste quality representation in the human brain

**DOI:** 10.1101/726711

**Authors:** Jason A. Avery, Alexander G. Liu, John E. Ingeholm, Cameron D. Riddell, Stephen J. Gotts, Alex Martin

## Abstract

In the mammalian brain, the insula is the primary cortical substrate involved in the perception of taste. Recent imaging studies in rodents have identified a gustotopic organization in the insula, whereby distinct insula regions are selectively responsive to one of the five basic tastes. However, numerous studies in monkeys have reported that gustatory cortical neurons are broadly-tuned to multiple tastes, and tastes are not represented in discrete spatial locations. Neuroimaging studies in humans have thus far been unable to discern between these two models, though this may be due to the relatively low spatial resolution employed in taste studies to date. In the present study, we examined the spatial representation of taste within the human brain using ultra-high resolution functional magnetic resonance imaging (MRI) at high magnetic field strength (7-Tesla). During scanning, participants tasted sweet, salty, sour and tasteless liquids, delivered via a custom-built MRI-compatible tastant-delivery system. Our univariate analyses revealed that all tastes (vs. tasteless) activated primary taste cortex within the bilateral dorsal mid-insula, but no brain region exhibited a consistent preference for any individual taste. However, our multivariate searchlight analyses were able to reliably decode the identity of distinct tastes within those mid-insula regions, as well as brain regions involved in affect and reward, such as the striatum, orbitofrontal cortex, and amygdala. These results suggest that taste quality is not represented topographically, but by a combinatorial spatial code, both within primary taste cortex as well as regions involved in processing the hedonic and aversive properties of taste.

## 1 INTRODUCTION

The sensation of taste begins on the tongue with sensory receptor cells tuned to one of the five basic tastes: sweet, salty, sour, bitter, and umami (Roper and Chaudhari, 2017). These taste receptor cells signal to afferent sensory neurons, and although some are dedicated to one specific taste, most receive input from multiple taste cells (Roper and Chaudhari, 2017). Thus taste coding in the periphery is a mixture of labelled-line transmission and a complex combinatorial code across afferent neurons that are both specifically and broadly-tuned (Simon et al., 2006; Roper and Chaudhari, 2017). In the taste afferent pathway, taste fibers travel along cranial nerves VII, IX, and X to reach their first synapse at the nucleus of the solitary tract in the medulla (Beckstead et al., 1980). In primates, these second-order taste afferents are relayed to the thalamic gustatory nucleus (knows as either the VMb or VPMpc nucleus) (Pritchard et al., 1986). At both of these sub-cortical relays different tastes are intermingled, with little evidence for topographic organization (Simon et al., 2006) prior to reaching their cortical target in the mid-dorsal region of the insular cortex (Small, 2010).

There are two basic models that can account for the representation of taste quality within the insular cortex: A *topographic* model, where a specific taste, such as sweet, is represented at a discreet spatial location, and a *combinatorial coding* model in which taste is represented by a combination of broadly tuned cortical neurons that are differentially sensitive to different tastes. The topographic model would be analogous to somatotopy in the somatosensory system, or retinotopy in the visual system. It should be noted, however, that unlike retinotopy and somatotopy, where sensory signals are represented according to their well-defined location in space, the sensation of taste has no such spatial layout, as previous theories of a taste map on the tongue have been disproven (Chandrashekar et al., 2006). Currently, although there is some support for the topographical model in rodents (Chen et al., 2011), human neuroimaging studies have failed to provide evidence for this possibility. These previous studies, however, have relied on small sample sizes and relatively low resolution imaging (Schoenfeld et al., 2004; Prinster et al., 2017). Higher resolution neuroimaging methods may be needed in order to identify to pographically distinct insula regions selective for particular tastes. Relatedly, another possibility suggested by a previous human neuroimaging study (Schoenfeld et al., 2004) is that, while distinct tastes may be represented topographically, this representation may not be uniform from one subject to another. If so, then a taste topography may not be observable at the group level but may be revealed in individuals scanned on different days.

In contrast to topographical models, many neurophysiology studies in rodents and primates have provided evidence for some form of a combinatorial coding model. These studies report that the neurons in gustatory cortex are responsive to a complex variety of orosensory, viscerosensory, and motor signals, with relatively few neurons responsive solely to taste (Scott and Plata-Salamn, 1999; Scott and Giza, 2000). Indeed, in primates, most taste-responsive neurons appear to be broadly tuned to multiple tastes, and there appears to be no observable topographic organization for specific tastes (Scott et al., 1991). If, as these primate studies suggest, (Scott and Plata-Salamn, 1999; Kaskan et al., 2018), insula gustatory regions are broadly responsive to multiple tastes, taste-specific neural responses might be discernable within those regions using multivariate pattern analysis (MVPA) techniques (Kriegeskorte et al., 2006). In fact, some evidence for this possibility in the human brain has recently been reported (Chikazoe et al., 2019).

To distinguish between these competing models, we examined taste-evoked hemodynamic responses within the human brain using ultra-high resolution functional magnetic resonance imaging (MRI) at high magnetic field strength (7-Tesla). During scanning, participants tasted sweet, salty, sour and tasteless (control) liquids, delivered via a custom-built MRI-compatible tastant-delivery system. To identify brain regions exhibiting shared and distinct responses for each taste, we employed both standard univariate analyses as well as multivariate analysis techniques to examine distributed activation patterns across multiple brain voxels.

## 2 METHODS

### 2.1 Participants

18 subjects (11 female) between the ages of 22 and 48 (Average: 27 years). Ethics approval for this study was granted by the NIH Combined Neuroscience Institutional Review Board under protocol number 10-M-0027. The institutional review board of the National Institutes of Health approved all procedures, and written informed consent was obtained for all subjects. Participants were excluded from taking part in the study if they had any history of neurological injury, known genetic or medical disorders that may impact the results of neuroimaging, prenatal drug exposure, severely premature birth or birth trauma, current usage of psychotropic medications, or any exclusion criteria for MRI.

### 2.2 Experimental Design

All scanning was performed at the NIH Clinical Center in Bethesda, MD. Participant sessions began with a taste assessment performed within the 7T scanner itself, but prior to the beginning of the scan session. This was done to ensure that ratings were obtained as close in time as possible to the scan itself, without any effects of differing environmental context between the testing and scanning session. The taste assessment was followed by a high resolution anatomical reference scan and a functional MRI session, during which they performed our Taste Perception task. Five participants returned for a second session where they once again completed the Taste Perception task. The time between these sessions varied from 6 to 98 days (average 34 days).

#### 2.2.1 Taste Stimuli

Participants received four tastant solutions during scanning and the pre-scan taste assessment: Sweet (0.6M sucrose), Sour (0.01M citric acid), Salty (0.20M NaCl), and Neutral (2.5mM NaHCO3 + 25mM KCl). To reduce within-subject variability, all participants received tastants at the same molar concentration. The specific concentrations used for this study were derived from median concentrations used in previous neuroimaging studies of taste perception, compiled in previous neuroimaging meta-analyses (Veldhuizen et al., 2011; Yeung et al., 2017). All tastants were prepared using sterile lab techniques and USP-grade ingredients by the NIH Clinical Center Pharmacy.

#### 2.2.2 Gustometer Description

A custom-built pneumatically-driven MRI-compatible system delivered tastants during fMRI-scanning (Simmons et al., 2013b; Avery et al., 2015; 2017; 2018), see Figure 1. Tastant solutions were kept at room temperature in pressurized syringes and fluid delivery was controlled by pneumatically-driven pinch-valves that released the solutions into polyurethane tubing that ran to a plastic gustatory manifold attached to the head coil. The tip of the polyethylene mouthpiece was small enough to be comfortably positioned between the subjects teeth. This insured that the tastants were always delivered similarly into the mouth. The pinch valves that released the fluids into the manifold were open and closed by pneumatic valves located in the scan room, which were connected to a stimulus delivery computer, which controlled the precise timing and quantity of tastants dispensed to the subject during the scan. Visual stimuli for behavioral and fMRI tasks were projected onto a screen located inside the scanner bore and viewed through a mirror system mounted on the head-coil. Both visual stimulus presentation and tastant delivery were controlled and synchronized via a custom-built program developed in the PsychoPy2 environment.

**Figure 1.**
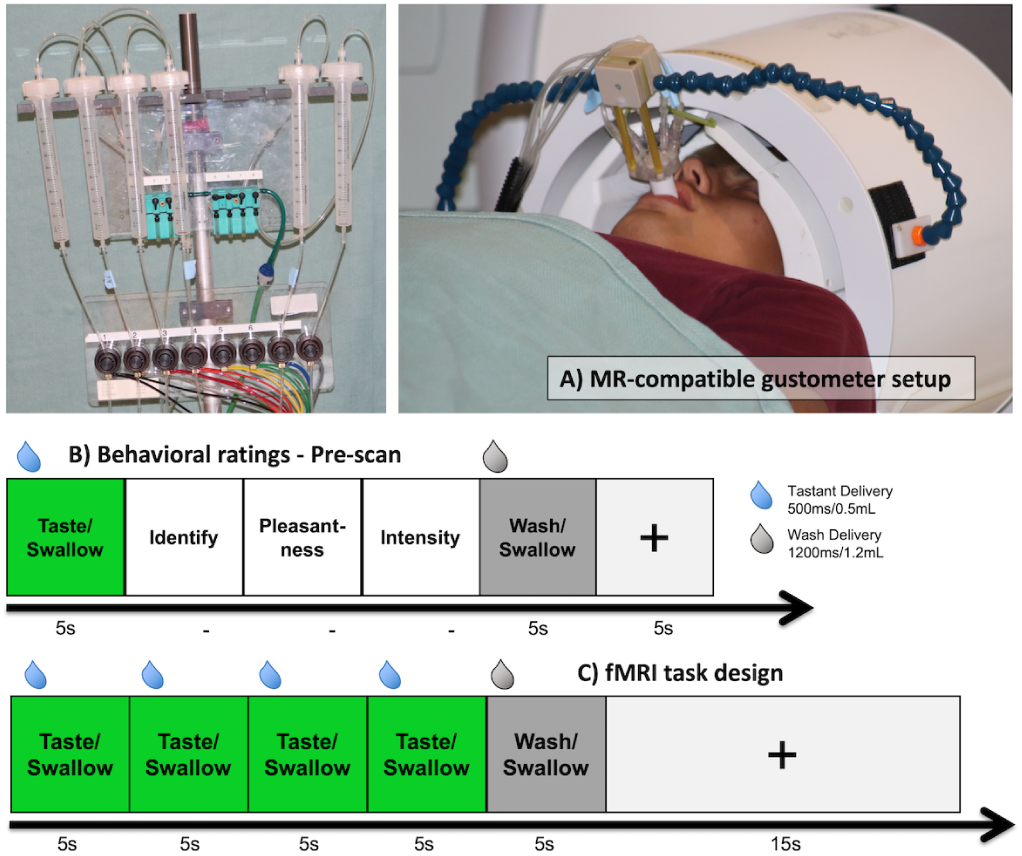
Tastant Delivery System and Task Design. A) During scanning, participants tasted sweet, salty, sour and tasteless liquids, delivered via a custom-built MRI-compatible tastant-delivery system. B) While in the scanner, but prior to fMRI scanning, subjects rated the identity, pleasantness, and intensity of each taste solution. During the fMRI task, received 0.5mL of sweet, sour, salty, and neutral tastant, in a block design (4 identical taste events/block), followed by a wash period.

#### 2.2.3 Taste Assessment

During blocks of the taste assessment task, the word Taste appeared on the screen for 2.5s, and subjects received 0.5mL of either a sweet, sour, salty, or neutral tastant (Figure 1b). Next, the word swallow appeared on the screen for 2.5s, prompting subjects to swallow the tastant. Following this, subjects were asked to indicate, using a hand-held response box, A) the identity of the tastant they received, by selecting it from one of four possible options (sweet, sour, salty, or neutral) presented on the screen B) the pleasantness of the taste on a 0 (not pleasant at all) to 10 (extremely pleasant) scale, and C) the intensity of the taste, on a 0 (not intense at all) to 10 (extremely intense) scale. Subjects provided ratings via a hand-held fiber optic response box (Current Designs, Philadelphia, PA). Rating periods were self-paced, average of 3s for rating trials and 2s for identification trials. Following these rating periods, the word wash appeared on the screen and subjects received 1.2mL of the neutral tastant, to rinse out the preceding taste. This was followed by another (2.5s) prompt to swallow. A five-second fixation period separated successive blocks of the taste assessment task. Tastants were presented five times each (20 blocks total), in random order. Altogether, this session lasted between five to ten minutes.

#### 2.2.4 Taste Perception task

During the Taste Perception fMRI task (Figure 1c), the word Taste appeared on the screen for 2.5s, and subjects received 0.5mL of either a sweet, sour, salty, or neutral tastant. Next, the word swallow appeared on the screen for 2.5s, prompting subjects to swallow. These taste and swallow periods occurred four times in a row, with the identical tastant delivered each time. Following these four peiods, the word wash appeared on the screen and subjects received 1.2mL of the neutral tastant, to rinse out the preceding tastes. This was followed by another (2.5s) prompt to swallow. In total, these taste delivery blocks lasted 25 seconds. Participants were not instructed to perform any ratings during the scanning session. Nor were they asked the identity, pleasantness, or intensity of the tastants. Previous research has identified that the insula is particularly sensitive to attentional orientation (Veldhuizen et al., 2007; Simmons et al., 2013a; Avery et al., 2015; 2017), and taste activity patterns vary significantly within the insula, according to task context (Bender et al., 2009). Following taste delivery blocks, a fixation cross + appeared on the screen for 15 seconds. Two sweet, salty, sour, and neutral taste delivery blocks (16 total) were presented in random order throughout each run of this task. Each run lasted 325 seconds (5min25sec). Participants completed 8 runs of the Taste Perception task during one scan session.

### 2.3 Imaging Methods

fMRI data was collected at the NIMH fMRI core facility at the NIH Clinical Center using a Siemens 7T-830/AS Magnetom scanner and a 32-channel head coil. Each volume consisted of 68 1.2-mm axial slices (echo time (TE) = 23 ms, repetition time (TR) = 2500 ms, flip angle = 55 degrees, voxel size = 1.2 1.2 1.2 mm3). A Multi-Band factor of 2 was used to acquire data from multiple slices simultaneously. A GRAPPA factor of 2 was used for in-plane slice acceleration along with a 6/8 partial Fourier k-space sampling. Each slice was oriented in the axial plane, with an Anterior-to-Posterior phase encoding direction. The scan window did not achieve full brain coverage though, and data was not acquired from areas at the top of the frontal and parietal lobes, as well as the very bottom of the cerebellum. See Figure 2a and 2c for boundaries of EPI scan window at subject and group level. Prior to task scans, a 1-minute EPI scan was acquired with the opposite phase encoding direction (Posterior-to-Anterior), which was used for correction of spatial distortion artifacts during preprocessing (see Image Preprocessing). An ultra-high resolution MP2RAGE sequence was used to provide an anatomical reference for the fMRI analysis (TE = 3.02 ms, TR = 6000 ms, flip angle = 5 degrees, voxel size = 0.70 0.70 0.70 mm).

**Figure 2.**
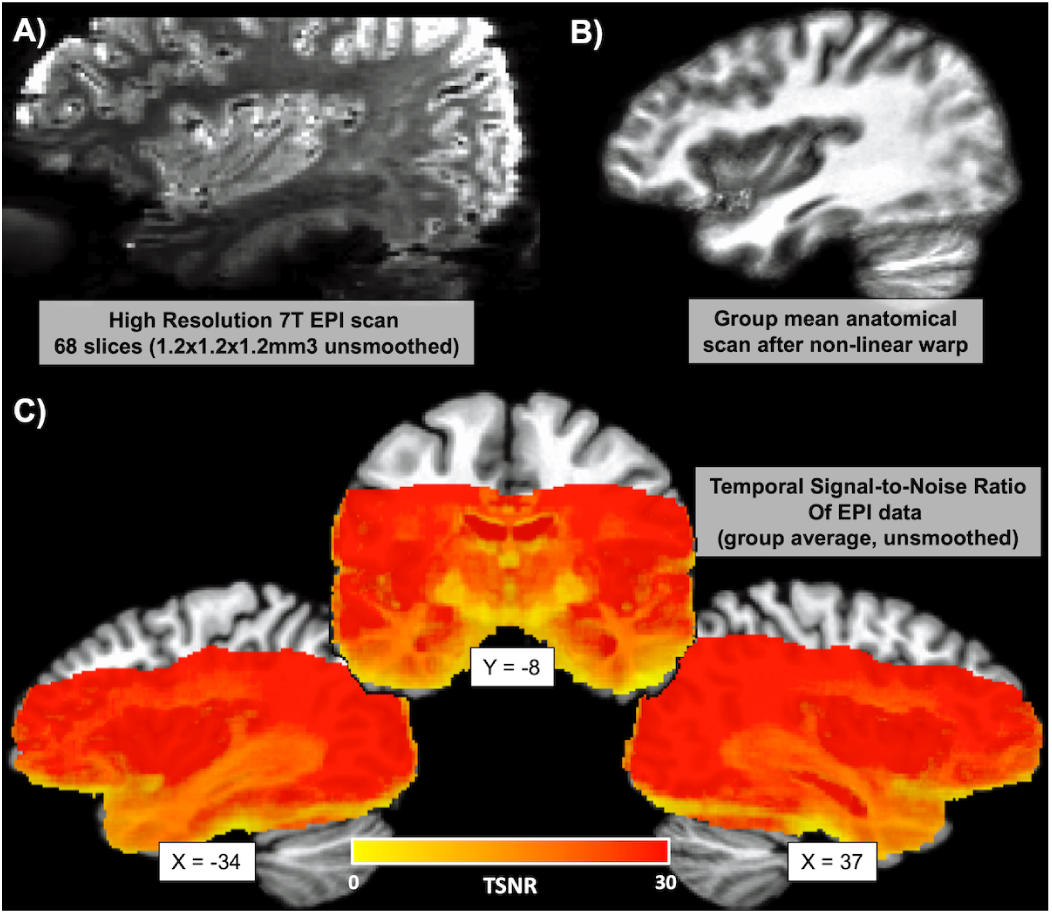
Imaging Parameters. Functional MRI data was acquired at ultra-high voxel resolution (1.2mm × 1.2mm × 1.2mm) at high magnetic field strength (7-Tesla). A) Echo-planar images (EPI) were acquired in 68 axial slices, in a scan window that ranged from the top of the cingulate gyrus (superiorly) to the tip of the temporal pole (inferiorly). B) Anatomical images were transformed to Talairach atlas space using a high-fidelity non-linear warp. C) Average temporal signal-to-noise ratio (TSNR) of unblurred EPI images, within a mask constructed from the intersection of EPI scan windows for all subjects, after non-linear transformation to Talairach space.

#### 2.3.1 Image Preprocessing

All fMRI pre-processing was performed in AFNI (http://afni.nimh.nih.gov/afni). The FreeSurfer software package (http://surfer.nmr.mgh.harvard.edu/) was additionally used for skull-stripping the anatomical scans. A de-spiking interpolation algorithm (AFNIs 3dDespike) was used to remove transient signal spikes from the EPI data, and a slice timing correction was then applied to the volumes of each EPI scan. The EPI scan acquired in the opposite (P-A) phase encoding direction was used to calculate a non-linear transformation matrix, which was used to correct for spatial distortion artifacts. All EPI volumes were then registered to the very first EPI volume using a 6-parameter (3 translations, 3 rotations) motion correction algorithm, and the motion estimates were saved for use as regressors in the subsequent statistical analyses. Volume registration and spatial distortion correction were implemented in the same non-linear transformation step, in order to minimize the number of interpolation steps performed on EPI data. For univariate analyses, smoothing with a 2.4mm (i.e. 2-voxel width) full width at half maximum Gaussian kernel was used to enhance image signal-to-noise ratio. Importantly, the core univariate results (see Figures 3 and 4) were not qualitatively different when examined with no spatial smoothing applied. No spatial smoothing was used for multivariate analyses. Finally, the signal intensity for each EPI volume was normalized to reflect percent signal change from each voxels mean intensity across the time-course.

**Figure 3.**
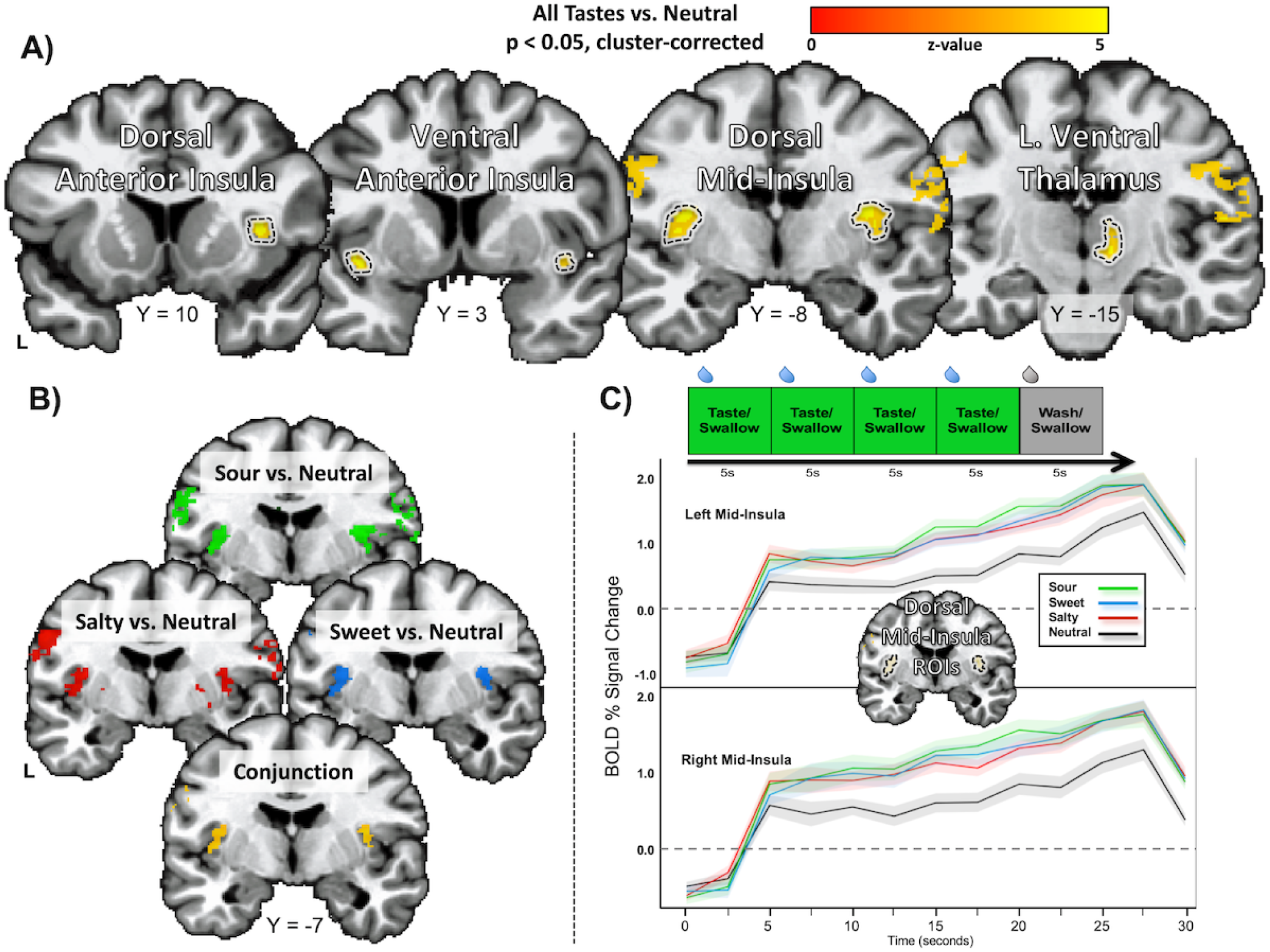
Taste-responsive regions of the brain. A) All tastes (sweet, sour, and salty) reliably activated taste cortex in the dorsal mid-insula, as well as primary lingual somatosensory cortex and a region of left ventral thalamus. Each of these regions has been previously identified as taste-responsive in prior human neuroimaging studies of taste (Small et al, 2010; Veldhuizen et al., 2011; Yeung et al., 2017). B) A conjunction of contrast maps for sweet, salty, and sour vs. neutral, separately corrected for multiple comparisons, identifies the bilateral dorsal mid-insula as commonly activated by all tastes. C) The hemodynamic response within the bilateral dorsal midinsula, estimated using finite impulse response functions, closely tracks the delivery of tastants during tastant blocks. The average response to each taste within the dorsal mid-insula is greater than the tasteless control, and this region exhibits no preferred activation for either taste.

**Figure 4.**
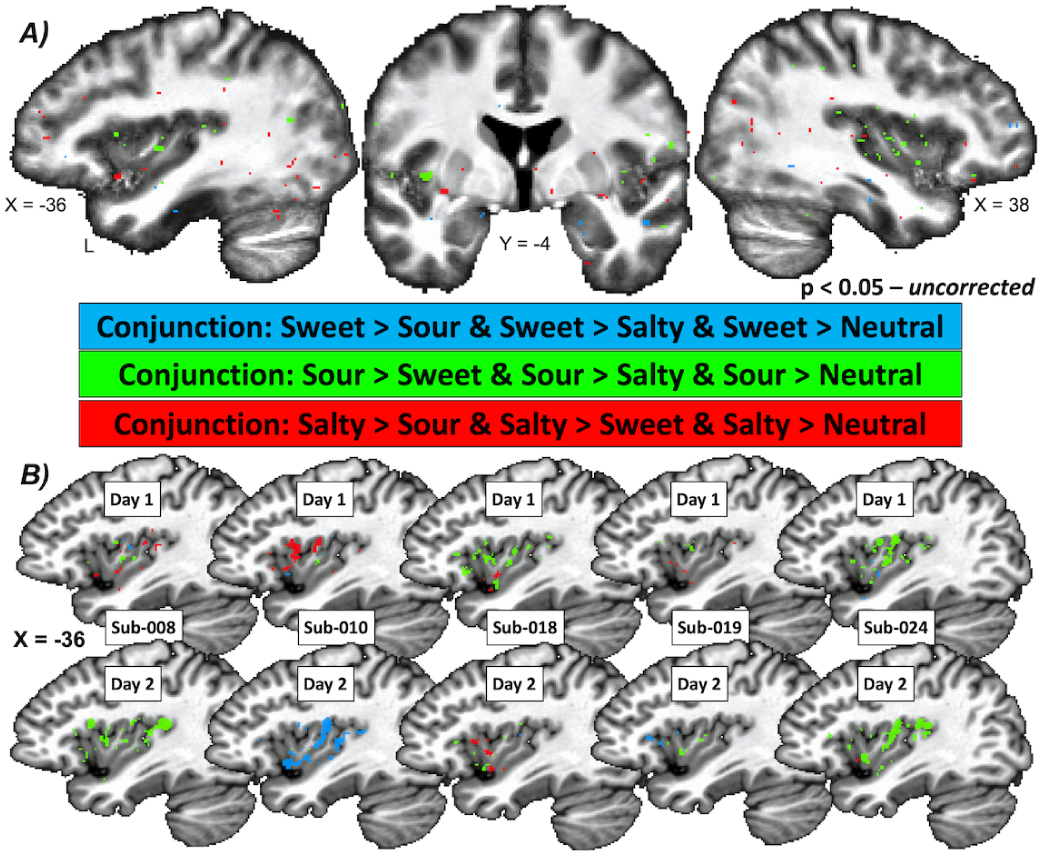
Conjunction analyses of taste contrasts provide no clear evidence of topographical organization. A) At the group level, whole brain conjunctions of taste contrasts, using corrected statistical contrasts, did not identify any voxels within the insula or the wider brain exhibiting a significant preference for either sweet, sour, or salty taste. B) Specificity maps from five subjects who were scanned on two separate sessions. Not only did specificity maps vary greatly between subjects, the patterns of taste-selective responses identified did not remain consistent within individual subjects from day to day. Images above show uncorrected specificity maps at low statistical threshold (p *<* 0.05).

Anatomical scans were first co-registered to the EPI scans and were then spatially normalized to Talairach space via a non-linear transformation implemented in AFNIs 3dQwarp. These non-linear transformation matrices were then used to register the subject-level statistical maps to Talairach atlas space.

In addition to volume registration, motion-censoring algorithms were also implemented to guard against potential artifactual confounds induced by uncontrolled subject motion. Briefly, the Euclidean-normalized derivative of the subjects motion parameters was calculated for each TR, and a list of time points was created in which that value was greater than 0.3 (roughly 0.3mm motion). The TRs within this list were then censored during the subject-level regression analysis. Additionally, any subject with an average Euclidean-normalized derivative of greater than 0.3 during any of the tasks was excluded from the group-level analysis. Three subjects were thus excluded from the original group of 21 subjects, due to excessive head motion, leaving 18 subjects.

The EPI data collected during the Taste Perception task were analyzed at the subject level using multiple linear regression models in AFNIs 3dDeconvolve. The regression model included one regressor for each tastant block (sweet, salty, sour, and neutral) and one regressor for wash/swallow events. These regressors were constructed by convolution of a gamma-variate hemodynamic response function with a boxcar function having a 20-second width (5 for wash/swallow events) beginning at the onset of each trial period. The regression model also included regressors of non-interest to account for each runs mean, linear, quadratic, and cubic signal trends, as well as the 6 normalized motion parameters (3 translations, 3 rotations) computed during the volume registration preprocessing.

### 2.4 Imaging Analyses

To evaluate between the competing models of taste representation, we performed both univariate and multivariate analyses on data from the Taste Perception task. For the univariate analyses, we sought to identify brain areas exhibiting both shared and distinct activation for each taste. We used the subject-level beta coefficient maps to perform group-level random-effects analyses, using the AFNI program 3dttest++. All group-level contrast maps created (sweet vs. sour, sweet vs. salty, salty vs. sour, and each taste vs. neutral) were separately corrected for multiple comparisons using non-parametric permutation tests, as described below.

Firstly, to identify brain areas exhibiting shared activation for each taste, we used the statistical contrast, all taste (sweet, salty, and sour) vs. neutral. Secondly, to identify brain voxels exhibiting specific taste preferences, we performed a series of conjunctions of our taste contrast maps. For each of our tastes (sweet, salty, and sour), we identified brain voxels where activity for that taste was greater than both other tastes, as well as the neutral tastant. For example, to be considered as sweet-selective the voxels would need to exhibit a significantly greater response to sweet versus sour, versus salty, and versus neutral. We performed these conjunction analyses at the level of individual subjects as well, separately correcting the statistical contrast maps that went into each conjunction using a voxel-wise FDR correction. In a less stringent approach to identifying differences between tastes at the group level, we also used the AFNI program 3dANOVA2 to identify a main effect of taste from among the three tastants (i.e. sweet, salty, and sour).

#### 2.4.1 Multivariate pattern analyses

Our next analysis approach was to identify whether, within insula regions broadly responsive to multiple tastes, taste-specific neural responses might be discernable using multivariate pattern analysis (MVPA) techniques. The analysis, based on a linear support-vector-machine approach, was trained and tested on subject-level regression coefficient maps obtained from the Taste Perception task, using leave-one-run-out cross-validation. These analyses were conducted using The Decoding Toolbox (Hebart et al., 2014). The insula regions-of-interest (ROIs) used for these analyses were created from the conjunction of sweet vs. neutral, sour vs. neutral, and salty vs. neutral contrasts, each separately corrected for multiple comparisons, as defined below.

At the whole-brain level, we ran a multivoxel pattern analysis (MVPA), using a searchlight approach (Kriegeskorte et al., 2006), which allowed us to identify the average classification accuracy of a multi-voxel searchlight centered on each voxel in the brain. The MVPA analyses were again conducted using The Decoding Toolbox (Hebart et al., 2014), and results were tested in AFNI. To generate these maps, a separate subject-level regression model was applied to the EPI data, which modeled each of the eight task runs separately, so that all task conditions would have eight beta-coefficient maps for the purposes of model training and testing. For every subject, we performed separate searchlight analyses for all three pairwise taste contrasts (sweet vs. sour, sweet vs. salty, salty vs. sour). The output of each searchlight analysis was a voxel-wise map of classification accuracy minus chance (50% for pairwise comparisons).

To evaluate the classification results at the group level, we averaged those three pairwise comparison maps together, warped the resulting map to Talairach atlas space, and applied a small amount (2.4mm FWHM) of spatial smoothing to normalize the distribution of scores across the dataset. We then performed group-level random-effects analyses using the AFNI program 3dttest++ and applied a non-parametric permutation test to correct for multiple comparisons.

To verify the specificity of those classification maps, we also performed separate group-level voxel-wise corrected t-tests for each of the pairwise taste contrasts and calculated the conjunction of all three of those corrected maps.

#### 2.4.2 Assessment of covariance with pleasantness ratings

We examined the relationship between fMRI activation to tastants and participants self-reported pleasantness ratings of those tastants in two separate ways. Firstly, we included participants average pleasantness ratings for each of the tastant solutions (including the neutral solution) in the subject-level regression model, using amplitude modulation regression in AFNI, which varies the height of the predicted hemodynamic response by the height of the associated behavioral covariate. In this case, the predicted response to tastant delivery varied as a function of subjects pleasantness ratings. The regression model also included regressors of non-interest, for motion, polynomial trend, and wash trials, as above. We then performed group-level random-effects analyses on the resulting amplitude-modulated beta-coefficient maps using the AFNI program 3dttest++. This first method allowed us to estimate the amount of within-session variability accounted for by participants pleasantness ratings.

Secondly, we performed a linear mixed-effects (LME) analysis at the group level, using the AFNI program 3dLME (Chen et al., 2013). For this analysis, we provided the program with the subject-level beta-coefficient maps for each tastant (including neutral), as well as subjects pleasantness ratings for each of those tastants. This analysis was used to estimate the amount of variability accounted for by pleasantness ratings, across subjects. This method also served as a means of removing the variance associated with that effect from our standard univariate contrasts (i.e. sweet vs. sour, sweet vs. salty, salty vs. neutral).

#### 2.4.3 Permutation testing for multiple comparison correction

Multiple comparison correction was performed using AFNIs 3dClustSim (AFNI version 19.0.25, March 15, 2019), within a whole-volume TSNR mask. This mask was constructed from the intersection of the EPI scan windows for all subjects (after transformation to talairach space) with a brain mask in atlas space (Figure 2c). The mask was then subjected to a TSNR threshold, such that all remaining voxels within the mask had an average un-smoothed TSNR of 10 or greater. For one-sample t-tests, this program will randomly flip the sign of individual datasets within a sample 10,000 times. This process generates an empirical distribution of clustersize at the desired voxel-wise statistical threshold (in this case, p *<* 0.001). The clusters which survive correction are those that are larger than 95% of the clusters within this empirical clustersize distribution. Permutation tests utilizing cluster size have been demonstrated to accurately control false positive rates (see (Eklund et al., 2016), for discussion).

## 3 RESULTS

### 3.1 Behavioral Results

Sweet, Salty, and Sour tastants did not differ in intensity (p *>* 0.10), but each was rated more intense than the neutral tastant (p *<* 0.01). The sweet tastant was rated as the most pleasant (p *<* 0.001), followed by the neutral tastant. The sour and sweet tastants did not differ in pleasantness (p = 0.83). Average identification accuracy was 93%(SD = 8%).

### 3.2 Imaging Results

#### 3.2.1 Univariate Contrasts

All tastes compared to the tasteless control significantly activated a bilateral region of the dorsal mid-insula, located in an area in the fundus of the central insular sulcus and the overlying frontoparietal operculum (Figure 3a, Table 1). In the left insula, as well as the right, to a lesser extent, activation for taste vs. neutral extended ventrally from this dorsal mid-insula area down the central insular sulcus, extending towards the ventral anterior insula. A separate cluster was located in the dorsal right anterior insula, located in the fundus of the overlying frontal operculum, on the middle short insular gyrus. Outside of the insula, we observed significant activation in the bilateral post-central gyrus, approximately in the tongue and mouth region of the motor strip, as well as bilateral regions of cerebellar lobule VI, regions associated with orosensory and motor movements (Suzuki et al., 2003). Finally, we observed activation for all tastes in a cluster of the right ventral thalamus, in the approximate location of the gustatory thalamic nucleus (Figure 3a, Table 1). These results were obtained using non-parametric multiple comparisons correction over the whole scan volume. In a follow up analysis, we performed a conjunction of contrast maps created by comparing each of the individual tastes vs. the tasteless control. This conjunction identified bilateral regions of the dorsal mid-insula and a region of the left somatosensory cortex (Figure 3b).

**Table 1.** Brain regions responsive to tastants (vs. tasteless control)

| Anatomical Location | Peak* | | | Peak $z$ | Cluster $p$ | Volume (mm<sup>3</sup>) |
| --- | --- | --- | --- | --- | --- | --- |
|  | X | Y | Z |  |  |  |
| R Somatosensory Cortex | +60 | -1 | +23 | 5.20 | < 0.01 | 2798 |
| L Somatosensory Cortex | -57 | -15 | +12 | 5.04 | < 0.01 | 2286 |
| L Cerebellum | -15 | -60 | -22 | 5.28 | < 0.01 | 1106 |
| L Mid-Insula | -34 | -7 | +14 | 5.46 | < 0.01 | 900 |
| R Cerebellum | +22 | -63 | -19 | 4.96 | < 0.01 | 757 |
| R Mid-Insula | +33 | -8 | +14 | 5.09 | < 0.01 | 689 |
| R Mid-Anterior Insula | +35 | +11 | +7 | 5.43 | < 0.01 | 313 |
| R Ventral Thalamus | +10 | -15 | +1 | 5.08 | < 0.01 | 257 |
| R Ventral Anterior Insula | +37 | +7 | -5 | 5.16 | < 0.02 | 152 |

\*Coordinates listed in Talairach and Tournoux stereotaxic space. L - Left; R - Right.

#### 3.2.2 Taste Specificity Analyses

A series of whole brain conjunction analyses was performed to identify voxels within the brain that were preferentially activated by each taste greater than both other tastes as well as the tasteless control. After correction for multiple comparisons and voxel-wise conjunction analysis, we did not identify any voxels within the insula or the wider brain exhibiting a significant preference for either sweet, sour, or salty taste. This procedure was repeated using multiple voxelwise statistical thresholds with identical results (Figure 4a, uncorrected maps presented for display purposes only). In another analysis approach, we did not identify any regions of the brain that exhibited a main effect of taste quality, after correction for multiple comparisons.

At the subject-level, a similar series of taste specificity analyses identified distributions of voxels within the insula that exhibited a selective preference for specific tastes. However, the distribution of these voxels was not consistent across subjects, nor were the dominant preferences in any consistent pattern (Figure 4b). To identify whether these taste-selective responses reflected a consistent spatial organization at the subject-level, we brought back a subset of our subjects for a second session of the Taste Perception task (N = 5). We observed that, within individual subjects, the patterns of taste-selective responses identified via the conjunction analyses did not remain consistent from day to day (Figure 4b).

#### 3.2.3 MVPA Results

To identify whether taste-selective activity patterns are present within insula regions broadly responsive to multiple tastes, we performed multivariate pattern analyses (MVPA) using the mid-insula ROIs identified above (Figure 3b). We were able to reliably classify between sweet, salty, and sour tastants within the bilateral dorsal mid-insula with an average accuracy of 60% (i.e. 10% above chance; all pairwise p-values p *<* 0.001; see Figure 5a).

**Figure 5.**
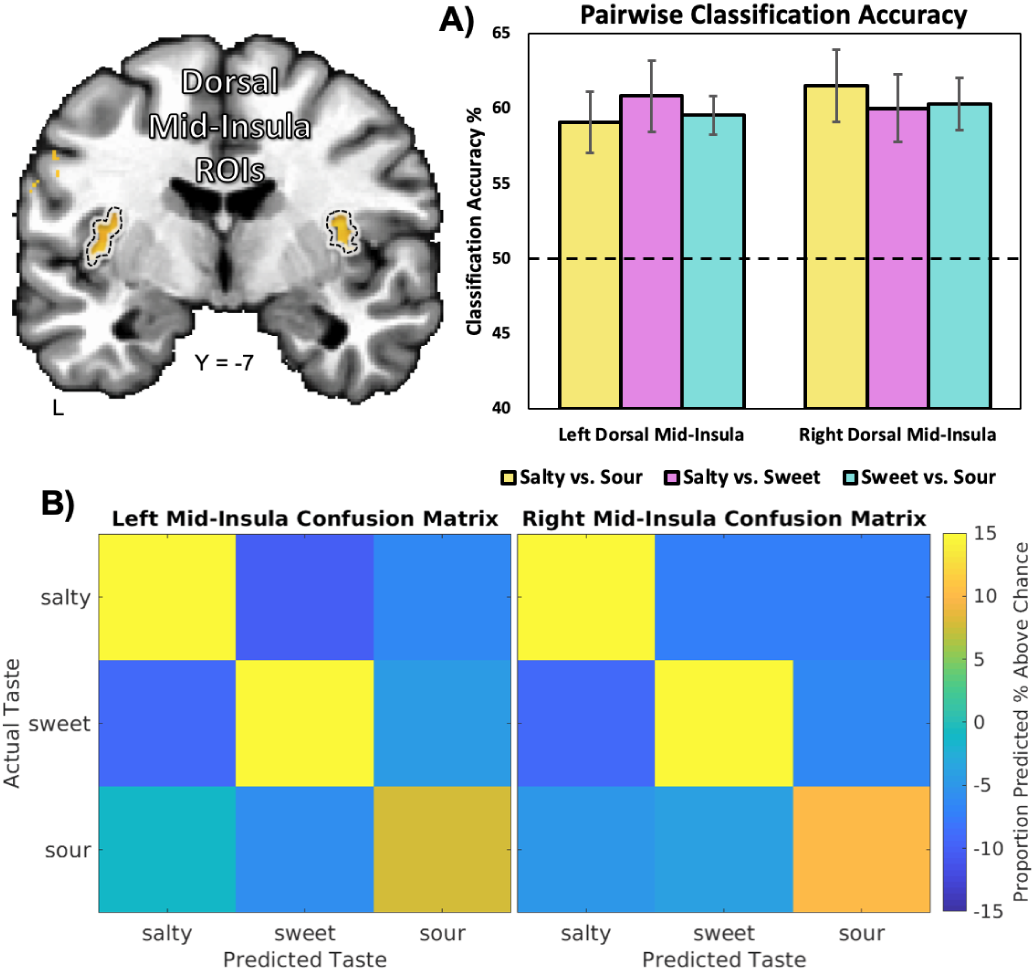
Multivariate pattern analyses decode taste quality within primary taste cortex. Multivariate pattern analyses run within the dorsal mid-insula, a region broadly responsive to multiple tastes (Figure 3b) show high accuracy for classifying between distinct tastes, as seen in A) pairwise classification accuracy plots and B) multi-class confusion matrices.

The multivariate searchlight analysis identified a number of brain regions that exhibited significant, above chance classification accuracy for discriminating between tastes. Those regions included primary sensory areas such as bilateral regions of the dorsal mid-insular cortex, somatosensory cortex, and piriform cortex. Additionally, we also were able to reliably classify between distinct tastes within brain regions involved in affect and reward, including the bilateral amygdala, the left orbitofrontal cortex (BA11m), the mediodorsal thalamus, the dorsal striatum, and the subgenual prefrontal cortex (BA25) (see Figure 6a and Table 2 for a list of these regions and descriptive and statistical data). In a complementary analysis, we performed a conjunction of the pairwise classification maps at the group level, each separately corrected for multiple comparisons. The conjunction analysis identified clusters within the bilateral mid-insula, left amygdala, and left piriform cortex (Figure 6b).

**Table 2.** Brain regions where multivoxel patterns reliably discriminate between sweet, salty, and sour tastants

| Anatomical Location | Peak* | | | Peak $z$ | Peak Acc.% | Cluster $p$ | Volume (mm<sup>3</sup>) |
| --- | --- | --- | --- | --- | --- | --- | --- |
|  | X | Y | Z |  |  |  |  |
| R Dorsal Mid-Insula | 36 | -9 | 10 | 5.65 | 14.00 | < 0.01 | 2964 |
| - R Parietal Operculum |  |  |  |  |  |  |  |
| R Piriform Cortex | 23 | -3 | -7 | 5.26 | 13.10 | < 0.01 | 1510 |
| L Amygdala | -20 | -4 | -13 | 5.47 | 13.90 | < 0.01 | 1337 |
| R Pons | 7 | -16 | -24 | 5.18 | 12.70 | < 0.01 | 1109 |
| L Somatosensory Cortex | -52 | -15 | 29 | 5.39 | 11.50 | < 0.01 | 1085 |
| L Dorsal Mid-Insula | -34 | -4 | 9 | 5.13 | 14.08 | < 0.01 | 931 |
| R Hippocampus | 19 | -23 | -12 | 4.86 | 11.50 | < 0.01 | 757 |
| R Somatosensory Cortex | 53 | -11 | 30 | 5.30 | 12.85 | < 0.01 | 755 |
| R Parahippocampal Gyrus | 28 | -27 | -19 | 4.80 | 11.80 | < 0.01 | 679 |
| L Periaqueductal Gray | -5 | -27 | -11 | 4.78 | 10.60 | < 0.01 | 662 |
| Subgenual Prefrontal Cortex | 8 | 25 | -3 | 5.05 | 13.40 | < 0.01 | 655 |
| L Cerebellum | -20 | -26 | -27 | 5.06 | 10.30 | < 0.01 | 534 |
| R Amygdala | 28 | 2 | -22 | 4.40 | 13.20 | < 0.01 | 491 |
| R Dorsal Lateral Thalamus | 14 | -10 | 9 | 5.26 | 11.20 | < 0.01 | 429 |
| L Temporal Pole | -27 | 11 | -29 | 5.35 | 13.80 | < 0.01 | 396 |
| L Lateral Orbitofrontal Cortex | -20 | 28 | -10 | 5.11 | 10.90 | < 0.01 | 359 |
| L Dorsal Striatum | -16 | 7 | 17 | 4.78 | 10.90 | < 0.01 | 223 |
| L Ventral Tegmental Area | -4 | -10 | -9 | 4.68 | 11.03 | < 0.02 | 219 |
| R Temporal Pole | 40 | 13 | -27 | 4.61 | 12.08 | < 0.02 | 207 |
| R Lingual Gyrus | 9 | -63 | 0 | 4.48 | 11.90 | < 0.02 | 200 |
| R Mediodorsal Thalamus | 4 | -17 | 6 | 5.29 | 9.93 | < 0.02 | 195 |
| R Hippocampus | 25 | -29 | -4 | 4.58 | 9.48 | < 0.02 | 188 |
| R Mid-Cingulate Cortex | 7 | 9 | 27 | 4.99 | 8.56 | < 0.03 | 175 |
| R Inferior Temporal Gyrus | 58 | -7 | -21 | 4.90 | 9.93 | < 0.03 | 173 |
| R Caudate Nucleus | 9 | 2 | 6 | 4.40 | 11.60 | < 0.03 | 173 |

\*Coordinates listed in Talairach and Tournoux stereotaxic space. L - Left; R - Right.

**Figure 6.**
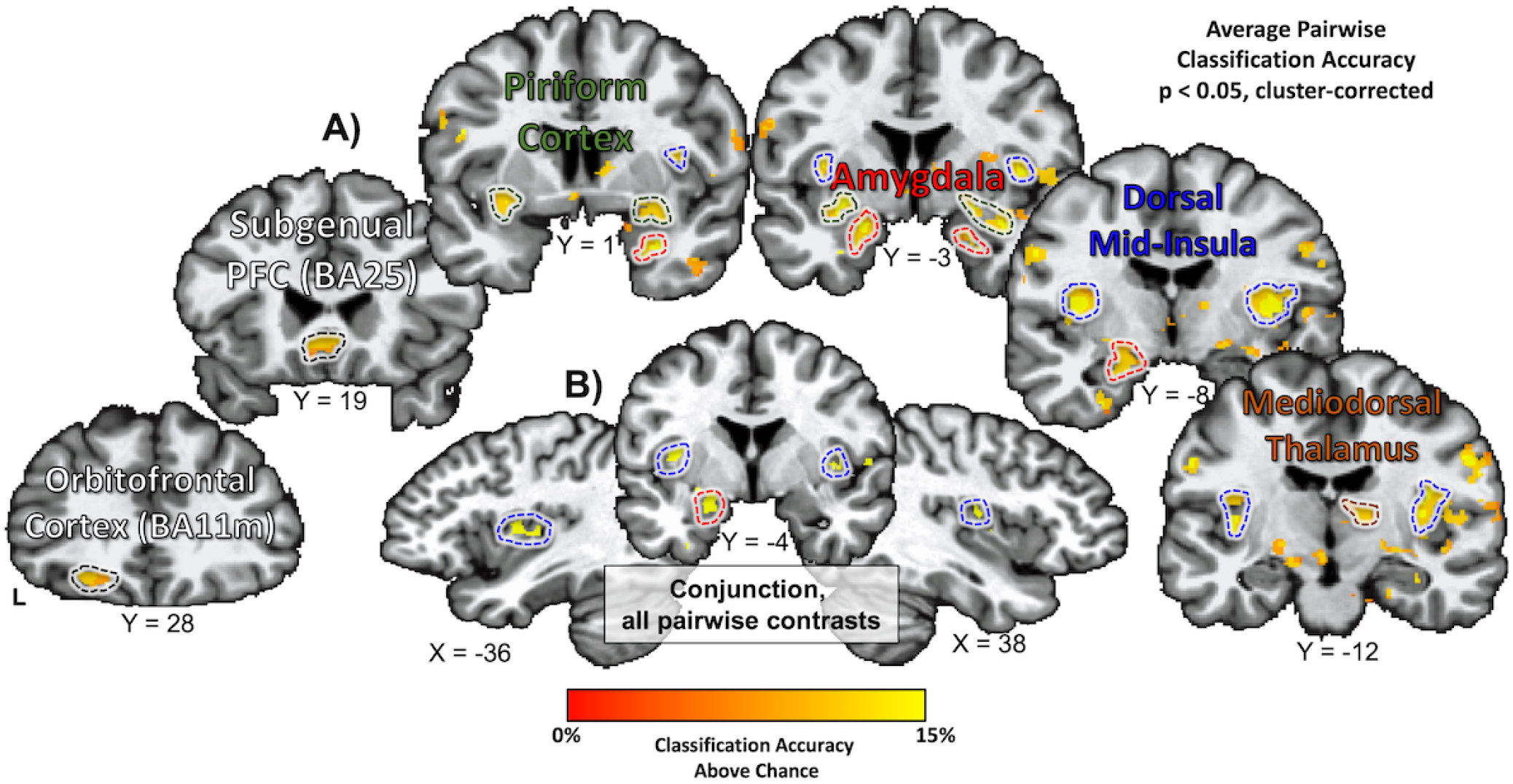
Multivariate pattern analyses decode taste quality within brain regions involved in taste perception and reward. (A) Several regions of the brain were identified using a multivariate searchlight analysis, trained to distinguish between sweet, salty, and sour tastes. Among those regions were primary sensory regions for taste the bilateral dorsal mid-insula and smell the bilateral piriform cortex, as well as limbic regions involved in affect and reward, such as the amygdala, orbitofrontal cortex, subgenual prefrontal cortex, and mediodorsal thalamus. B) A conjunction of separate searchlight analyses performed on pairwise taste comparisons (as in Figure 5a, above) identifies the bilateral mid-insula, left amygdala, and left piriform cortex (not pictured).

#### 3.2.4 Pleasantness Analyses

We also attempted to identify the effect of self-reported pleasantness ratings on the hemodynamic response to tastants. We performed both a group-level t-test of parametrically-modulated hemodynamic response functions as well as a group-level linear mixed-effects regression model. Neither approach was able to identify any brain regions exhibiting a reliable relationship between pleasantness ratings and tastant response. Additionally, removal of the effect of pleasantness ratings, using the LME model, did not significantly diminish the size of clusters present with our main univariate contrast (i.e. taste vs. tasteless, Figure 3).

## 4 DISCUSSION

Prior evidence from optical imaging studies in rodent models has suggested that tastes are represented in topographically distinct areas within the insular cortex (Chen et al., 2011). However, studies in non-human primates instead have suggested that taste within the insula is represented within populations of broadly tuned neurons, without any clear spatial organization (Scott et al., 1999). Human neuroimaging studies of taste to date have largely been conducted using low-resolution functional neuroimaging methods, which may not have the power to discriminate between distinct taste-specific regions within insular taste cortex. In order to discriminate between the competing topographical and combinatorial coding models, we examined the spatial representation of multiple tastes within the human brain using ultra-high resolution fMRI at high magnetic field strength. In agreement with previous human neuroimaging studies of taste (Small, 2010; Veldhuizen et al., 2011; Yeung et al., 2017), sweet, sour, and salty tastant solutions (vs. a tasteless solution) activated regions of the anterior and mid-insular cortex. However, we did not observe any evidence that any region within the insula, or in the rest of the brain, exhibited a clear preference for a specific taste.

One previous fMRI study that examined taste representation in the insula reported that patterns of taste-specific responses were highly variable across subjects, but suggested that those patterns were stable over time within subjects (Schoenfeld et al., 2004). To examine this possibility of a stable individual-level topography, we also examined the tastant responses of a subset of subjects which were scanned during two separate sessions. Using the same type of conjunction analyses we employed at the group level, we observed that the patterns of taste-specific responses were not only highly variable across subjects, but within subjects as well. While we were unable to rule out the possibility of a micro-scale organization of taste specific neurons within gustatory cortex, such as that observed in previous studies of rodents (Chen et al., 2011), we find little evidence for discrete topographical regions for specific tastes in humans at the spatial scales observable with ultra-high resolution fMRI.

However, using multivariate pattern analysis at those same spatial scales, we were nonetheless able to discriminate between the responses to distinct tastes within the insula. Using an MVPA searchlight approach (Kriegeskorte et al., 2006), we identified that the bilateral dorsal mid-insula reliably classified between distinct tastes with an accuracy significantly greater than chance. These results support a model of insular functional organization wherein taste quality is encoded by distinct spatial patterns within primary gustatory regions of the insular cortex (Simon et al., 2006). These activity patterns may be represented either in heterogenous pools of neurons specialized for detecting specific tastes, by a combinatorial population code within cortical neurons broadly tuned to multiple tastes, or by some mixture of broadly-tuned and specialist neurons, much like how taste is represented in the peripheral nervous system (Roper and Chaudhari, 2017).

We also explored the possibility that any effect of taste response that we observed was simply due to differences in relative pleasantness of tastant solutions. We tested this possibility using two different approaches, one in which participants pleasantness ratings were used to account for trial-by-trial variance (i.e.amplitude modulation regression) and one in which ratings were used to account for any remaining variance at the group level. Neither approach provided evidence for an effect of participants’ self-reported pleasantness ratings upon the hemodynamic response to tastes. Thus, within the parameters of this study, in which tastes were only mildly pleasant or mildly aversive, variability in pleasantness did not account for a variability in taste response.

The actual location of the human primary gustatory area in the insula has also been the subject of some controversy. Many researchers have argued that the far anterior region of the dorsal insula is the likeliest candidate, given the location of Area G in the non-human primate insula (Rolls, 2016). However, numerous human neuroimaging studies over the past 15 years have implicated the dorsal anterior insula as playing a more domain-general role in cognition and attention (Kurth et al., 2010; Nelson et al., 2010), and many consider the human anterior insula to be a newly developed cortical structure (Craig, 2009), perhaps due to the expansion of language faculties in our species (Nieuwenhuys, 2012). The effects of task context, such as whether subjects identify or evaluate a taste during scanning, have also been shown to affect the spatial representation of that taste within the insula (Bender et al., 2009). Nevertheless, a large number of human neuroimaging studies of taste, incorporating multiple imaging modalities (Kobayakawa et al., 1999; 2005; Veldhuizen et al., 2007), as well as cortical electrode stimulation studies in pre-surgical patients (Mazzola et al., 2017), point to the mid-to-posterior dorsal insula as the location of human primary gustatory cortex. In the present study, we identified regions of the dorsal mid-insula, anterior ventral insula, and mid-anterior insula that exhibited a significant activation to taste vs. tasteless solutions, consistent with meta-analyses of human neuroimaging studies (Veldhuizen et al., 2011; Yeung et al., 2017). Of those regions, the bilateral dorsal mid-insula regions exhibited the most consistent ability to discriminate between different tastant solutions, suggesting a focal role for the dorsal mid-insula in the primary sensory processing of taste.

One recent human neuroimaging study, which was acquired using lower resolution fMRI (36.75mm3 at 3T; 8mm3 at 7T vs. 1.73mm3 for the present study) and within the context of a hedonic evaluation task, also provided evidence that multivariate methods could be used to classify tastes within the insula (Chikazoe et al., 2019). In agreement with the results of Chikazoe and colleagues, we identified regions of the mid-insula that reliably classify between different tastes. In contrast to their results, however, we identified other regions of the brain that exhibited spatial activity patterns that discriminated between different tastes. These regions included other cortical sensory regions such as the piriform cortex, involved in olfactory processing (Sobel et al., 2000) as well as regions involved in various aspects of food sensation and reward, such as the bilateral amygdala, the orbitofrontal cortex, the dorsal striatum, and the mediodorsal thalamus. These results suggest that the presence of multivariate patterns supporting taste-specific information is not a unique feature of insular taste cortex, as previously suggested (Chikazoe et al., 2019).

Notably, the amygdala, orbitofrontal cortex and striatum are all downstream regions in the taste pathway (Scott and Plata-Salamn, 1999; Rolls, 2005), that receive primary (or secondary) projections from gustatory cortex, and that play different roles in food reward, aversion, and value based decision making (Kringelbach et al., 2003; Kringelbach, 2005; Rolls, 2005; Saez et al., 2017). The involvement of the amygdala in the perception of basic tastes has been demonstrated in patients undergoing amygdala resection, who exhibit increased sensitivity to sour taste and greater perceived taste intensity (Small et al., 1997). The activity of the amygdala, especially the central nucleus and basolateral amygdala, is strongly associated with conditioned taste aversion (Reilly and Bornovalova, 2005). The OFC, in concert with the amygdala and mediodorsal thalamus, are thought to represent the moment-to-moment value of environment stimuli and sensory experiences, informed by the bodys current state (Rudebeck and Murray, 2014). As these regions play key roles in taste perception and normative responses to food, the multivariate taste-specific patterns identified within them may reflect information related to the appetitive or aversive components of these tastant solutions.

## 4.1 Conclusion

We set out to distinguish between two competing theories of taste representation in the insula using ultra-high resolution fMRI at high magnetic field strength. We identified a strong overlap in the activity for all three tastes in previously identified gustatory regions of the insula. We did not identify any regions within the insula or the wider brain that exhibited a preference for specific tastes. However, we were able to decode taste identity in a consistent manner within primary gustatory insula and other brain regions associated with food perception and reward. This suggests that taste quality exists in a distributed pattern across multiple voxels within gustatory cortex, potentially resulting from a combinatorial code among populations of broadly tuned cortical neurons. This information is then presumably passed down through a network of cortical and sub-cortical regions involved in appetitive and behavioral responses to food.

## ACKNOWLEDGMENTS

The authors would like to thank Sean Marrett, Martin Hebart, the NIMH Section on Instrumentation and the NIH Clinical Center pharmacy for their assistance with various aspects of the design and execution of this study. This study was supported by the Intramural Research Program of the National Institute of Mental Health, National Institutes of Health, and it was conducted under NIH Clinical Study Protocol 10-M-0027 (ZIA MH002920). Clinicaltrials.gov ID: NCT01031407.

## REFERENCES

Avery JA, Gotts SJ, Kerr KL, Burrows K, Ingeholm JE, Bodurka J, Martin A, Simmons WK (2017) Convergent gustatory and viscerosensory processing in the human dorsal mid-insula. Hum brain mapp 38:2150–2164.

Avery JA, Ingeholm JE, Wohltjen S, Collins M, Riddell CD, Gotts SJ, Kenworthy L, Wallace GL, Simmons WK, Martin A (2018) Neural correlates of taste reactivity in autism spectrum disorder. NeuroImage: Clinical 19:38–46.

Avery JA, Kerr KL, Ingeholm JE, Burrows K, Bodurka J, Simmons WK (2015) A common gustatory and interoceptive representation in the human mid-insula. Hum brain mapp 36:2996–3006.

Beckstead RM, Morse JR, Norgren R (1980) The nucleus of the solitary tract in the monkey: projections to the thalamus and brain stem nuclei. J Comp Neurol 190:259–282.

Bender G, Veldhuizen MG, Meltzer JA, Gitelman DR, Small DM (2009) Neural correlates of evaluative compared with passive tasting. Eur J Neurosci 30:327–338.

Chandrashekar J, Hoon M, Ryba N (2006) The receptors and cells for mammalian taste. Nature 444:288–294.

Chen G, Saad ZS, Britton JC, Pine DS, Cox RW (2013) Linear mixed-effects modeling approach to FMRI group analysis. NeuroImage 73:176–190.

Chen X, Gabitto M, Peng Y, Ryba NJP, Zuker CS (2011) A gustotopic map of taste qualities in the mammalian brain. Science 333:1262–1266.

Chikazoe J, Lee DH, Kriegeskorte N, Anderson AK (2019) Distinct representations of basic taste qualities in human gustatory cortex. Nature Communications 10(1):1048.

Craig ADB (2009) How do you feel–now? The anterior insula and human awareness. Nat Rev Neurosci 10:59–70.

Eklund A, Nichols TE, Knutsson H (2016) Cluster failure: Why fMRI inferences for spatial extent have inflated false-positive rates. Proc Natl Acad Sci USA 113:201602413–201607905.

Hebart MN, Grgen K, Haynes J-D (2014) The Decoding Toolbox (TDT): a versatile software package for multivariate analyses of functional imaging data. Front Neuroinform 8:88.

Kaskan PM, Dean AM, Nicholas MA, Mitz AR, Murray EA (2018) Gustatory responses in macaque monkeys revealed with fMRI: Comments on taste, taste preference, and internal state. NeuroImage 184:932–942.

Kobayakawa T, Ogawa H, Kaneda H, Ayabe-Kanamura S, Endo H, Saito S (1999) Spatio-temporal analysis of cortical activity evoked by gustatory stimulation in humans. Chem Senses 24:201–209.

Kobayakawa T, Wakita M, Saito S, Gotow N, Sakai N, Ogawa H (2005) Location of the primary gustatory area in humans and its properties, studied by magnetoencephalography. Chem Senses 30 Suppl 1:i226–i227.

Kriegeskorte N, Goebel R, Bandettini P (2006) Information-based functional brain mapping. Proc Natl Acad Sci USA 103:3863–3868.

Kringelbach ML (2005) The human orbitofrontal cortex: linking reward to hedonic experience. Nat Rev Neurosci 6:691–702.

Kringelbach ML, O’Doherty J, Rolls ET, Andrews C (2003) Activation of the human orbitofrontal cortex to a liquid food stimulus is correlated with its subjective pleasantness. Cereb Cortex 13:1064–1071.

Kurth F, Zilles K, Fox PT, Laird AR, Eickhoff SB (2010) A link between the systems: functional differentiation and integration within the human insula revealed by meta-analysis. Brain Struct Funct 214:519–534.

Mazzola L, Mauguire F, Isnard J (2017) Electrical Stimulations of the Human Insula. Journal of Clinical Neurophysiology 34:307–314.

Nelson SM, Dosenbach NUF, Cohen AL, Wheeler ME, Schlaggar BL, Petersen SE (2010) Role of the anterior insula in task-level control and focal attention. Brain Struct Funct 214:669–680.

Nieuwenhuys R (2012) The insular cortex: a review. Prog Brain Res 195:123–163.

Prinster A, Cantone E, Verlezza V, Magliulo M, Sarnelli G, Iengo M, Cuomo R, Di Salle F, Esposito F (2017) Cortical representation of different taste modalities on the gustatory cortex: A pilot study. PLoS ONE 12:e0190164.

Pritchard TC, Hamilton RB, Morse JR, Norgren R (1986) Projections of thalamic gustatory and lingual areas in the monkey, Macaca fascicularis. J Comp Neurol 244:213–228.

Reilly S, Bornovalova MA (2005) Conditioned taste aversion and amygdala lesions in the rat: a critical review. Neurosci Biobehav Rev 29:1067–1088.

Rolls E (2005) Taste, olfactory, and food texture processing in the brain, and the control of food intake. Physiology & Behavior 85:45–56.

Rolls ET (2016) Functions of the anterior insula in taste, autonomic, and related functions. Brain and cognition 110:4–19

Roper SD, Chaudhari N (2017) Taste buds: cells, signals and synapses. Nat Rev Neurosci 18:485–497.

Rudebeck PH, Murray EA (2014) The Orbitofrontal Oracle: Cortical Mechanisms for the Prediction and Evaluation of Specific Behavioral Outcomes. Neuron 84:1143–1156.

Saez RA, Saez A, Paton JJ, Lau B, Salzman CD (2017) Distinct Roles for the Amygdala and Orbitofrontal Cortex in Representing the Relative Amount of Expected Reward. Neuron 95:70–77.e73.

Schoenfeld MA, Neuer G, Tempelmann C, Schler K, Noesselt T, Hopf JM, Heinze HJ (2004) Functional magnetic resonance tomography correlates of taste perception in the human primary taste cortex. Neuroscience 127:347–353.

Scott TR, Giza BK (2000) Issues of gustatory neural coding: where they stand today. Physiology & Behavior 69:65–76.

Scott TR, Plata-Salamn CR (1999) Taste in the monkey cortex. Physiology & Behavior 67:489–511.

Scott TR, Plata-Salamn CR, Smith VL, Giza BK (1991) Gustatory neural coding in the monkey cortex: stimulus intensity. J Neurophysiol 65:76–86.

Simmons WK, Avery JA, Barcalow JC, Bodurka J, Drevets WC, Bellgowan P (2013a) Keeping the body in mind: insula functional organization and functional connectivity integrate interoceptive, exteroceptive, and emotional awareness. Hum brain mapp 34:2944–2958.

Simmons WK, Rapuano KM, Kallman SJ, Ingeholm JE, Miller B, Gotts SJ, Avery JA, Hall KD, Martin A (2013b) Category-specific integration of homeostatic signals in caudal but not rostral human insula. Nat Neurosci 16:1551–1552.

Simon SA, De Araujo IE, Gutierrez R, Nicolelis MAL (2006) The neural mechanisms of gustation: a distributed processing code. Nat Rev Neurosci 7:890–901.

Small DM (2010) Taste representation in the human insula. Brain Struct Funct 214:551–561.

Small DM, Jones-Gotman M, Zatorre RJ, Petrides M, Evans AC (1997) A role for the right anterior temporal lobe in taste quality recognition. J Neurosci 17:5136–5142.

Sobel N, Prabhakaran V, Zhao Z, Desmond JE, Glover GH, Sullivan EV, Gabrieli JD (2000) Time course of odorant-induced activation in the human primary olfactory cortex. J Neurophysiol 83:537–551.

Suzuki M, Asada Y, Ito J, Hayashi K, Inoue H, Kitano H (2003) Activation of cerebellum and basal ganglia on volitional swallowing detected by functional magnetic resonance imaging. Dysphagia 18:71–77.

Veldhuizen MG, Albrecht J, Zelano C, Boesveldt S, Breslin P, Lundstrm JN (2011) Identification of human gustatory cortex by activation likelihood estimation. Hum brain mapp 32:2256–2266.

Veldhuizen MG, Bender G, Constable RT, Small DM (2007) Trying to detect taste in a tasteless solution: modulation of early gustatory cortex by attention to taste. Chem Senses 32:569–581.

Yeung AWK, Goto TK, Leung WK (2017) Basic taste processing recruits bilateral anteroventral and middle dorsal insulae: An activation likelihood estimation meta-analysis of fMRI studies. Brain and Behavior 7:e00655–12.

